# Highly Predictive Transdiagnostic Features Shared across Schizophrenia, Bipolar Disorder, and ADHD Identified Using a Machine Learning Based Approach

**DOI:** 10.1101/453951

**Authors:** Yuelu Liu, Monika S. Mellem, Humberto Gonzalez, Matthew Kollada, Atul R. Mahableshwarkar, Annette Madrid, William J. Martin, Parvez Ahammad

## Abstract

The Diagnostic and Statistical Manual of Mental Disorders (DSM) is the standard for diagnosing psychiatric disorders in the United States. However, evidence has suggested that symptoms in psychiatric disorders are not restricted to the boundaries between DSM categories, implying an underlying latent transdiagnostic structure of psychopathology. Here, we applied an importance-guided machine learning technique for model selection to item-level data from self-reported instruments contained within the Consortium for Neuropsychiatric Phenomics dataset. From 578 questionnaire items, we identified a set of features which consisted of 85 items that were shared across diagnoses of schizophrenia (SCZ), bipolar disorder (BD), and attention deficit/hyperactivity disorder (ADHD). A classifier trained on the transdiagnostic features reliably distinguished the patient group as a whole from healthy controls (classification AUC = 0.95) and only 10 items were needed to attain the performance level of AUC being 0.90. A sum score created from the items produced high separability between patients and healthy controls (Cohen’s d = 2.85), and it outperformed predefined sum scores and sub-scores within the instruments (Cohen’s d ranging between 0.13 and 1.21). The transdiagnostic features comprised both symptom domains (e.g. dysregulated mood, attention deficit, and anhedonia) and personality traits (e.g. neuroticism, impulsivity, and extraversion). Moreover, by comparing the features that were common across the three patient groups with those that were most predictive of a single patient category, we can describe the unique features for each patient group superimposed on the transdiagnostic feature structure. Overall, our results reveal a latent transdiagnostic symptom/behavioral phenotypic structure shared across SCZ, BD, and ADHD and present a new perspective to understand insights offered by self-report psychiatric instruments.

## 1 Introduction

The Diagnostic and Statistical Manual of Mental Disorders (DSM) provides a symptom-based taxonomy which serves to help clinicians classify various clusters of symptoms and abnormal behaviors into distinct categories of disorders. The uniformity of diagnostic criteria in DSM serves to effectively index psychiatric disorders but does not provide a data-driven framework within which to understand the shared and unique features across disorders. For example, dimensionality and comorbidity are pervasive in terms of symptoms across different DSM categories (Kessler et al., 2005; Markon, 2009; Krueger and Markon, 2011). Such dimensionality manifests as heterogeneity in symptom clusters within disease categories defined by the DSM and is exemplified across DSM categories (Kessler et al., 2007). In the area of anxiety and mood disorders, more than 50% of individuals are diagnosed as having more than a single category of disorders according to the DSM at a given time (Grisanzio et al., 2017). Similarly, about 50% of bipolar disorder patients exhibit schizophrenia-like psychotic symptoms during illness episodes (Coryell et al., 2001; Keck et al., 2003). The presence of such psychotic symptoms can be mood-incongruent (Pacheco et al., 2010) and can occur outside of illness episodes (Pope and Lipinski, 1978; Abrams and Taylor, 1981). These observations highlight the likelihood of a latent trans-diagnostic dimensional structure that spans multiple disorders (Krueger and Markon, 2006) and underscore the importance of understanding patients at the symptom-level, rather than simply at a diagnostic level, to create more effective treatments.

Studies have attempted to uncover the latent structure of psychopathology, between or within categories, through multimodal assessments that measure symptoms, behavior, physiology, imaging, and genetics. One such example is the large-scale study conducted by the UCLA Consortium for Neuropsychiatric Phenomics (CNP), which seeks to identify links among phenotypic data, imaging, and genetics (Poldrack et al., 2016). Overall, genetic studies have pointed to the heritability of major neuropsychiatric disorders (Cross-Disorder Group of the Psychiatric Genomics Consortium, 2013; Hamshere et al., 2013; Larsson et al., 2013; The Brainstorm Consortium et al., 2017; Bipolar Disorder and Schizophrenia Working Group of the Psychiatric Genomics Consortium et al., 2018; Gandal et al., 2018) as well as the genetic commonality amongst disorders (Purcell et al., 2009; Lotan et al., 2014) such as schizophrenia (SCZ), bipolar disorder (BD), and attention deficit/hyperactivity disorder (ADHD). Recent data-driven studies based on symptom and behavior have focused on classifying and subtyping patients within a single diagnostic category (Lamers et al., 2012; van Loo et al., 2012; Georgiades et al., 2013; Doshi-Velez et al., 2014; van Hulst et al., 2014; Costa Dias et al., 2015; Geisler et al., 2015; Sun et al., 2015; Drysdale et al., 2016; Gheiratmand et al., 2017). Several of these studies identified important shared abnormal features associated with the latent transdiagnostic structure across major psychiatric disorders.

Despite recent advancements, several unresolved issues still remain in the field. First, the clinical utility of using the features identified in the above-mentioned studies to reliably classify patients remains an open question. Emerging studies have used unsupervised machine learning approaches, such as clustering and dimensionality reduction algorithms, to uncover the transdiagnostic structure across disorders (Grisanzio et al., 2017; Xia et al., 2018). However, the lack of ground truth on how patients should be assigned to an identified cluster/subtype limits the application of these insights. Second, despite recent genetic studies documenting shared risk factors among SCZ, BD, and ADHD (Cross-Disorder Group of the Psychiatric Genomics Consortium, 2013; Larsson et al., 2013), a trans-diagnostic dimensional structure shared across the three disorders has not been discovered in other modalities such as neuroimaging and clinical characteristics. While a substantial body of neuroimaging studies have focused on investigating shared etiology between SCZ and BD (see e.g., Rashid et al., 2014), studies that further incorporated ADHD are scarce. Third, how well different feature modalities can be used as markers to reliably identify psychiatric patients in a clinical setting has not been systematically compared in prior literature. Studies typically reported systematic deviations within a single feature modality among psychiatric patients (Buckholtz and Meyer-Lindenberg, 2012; Goodkind et al., 2015; Sha et al., 2018) and the relative predictive power of various feature modalities in transdiagnostic scenarios remains unknown.

In the current study, we addressed the above issues by taking a patient-focused approach to identify transdiagnostic features that are shared across SCZ, BD, and ADHD. Following the definition used in the recent literature, we used the term “transdiagnostic” to denote features that extend beyond a single DSM category. Using an importance-guided model selection approach, the supervised machine learning framework used in this study allowed us to evaluate the performance of the transdiagnostic features and hence to iteratively identify the optimal set of features required to distinguish the patient group from healthy controls (HCs). Based on the CNP dataset, we used multiple data modalities including the behavioral/symptom phenotypes (from here on referred to as phenotypes) defined in self-reported instruments and neuroimaging data (sMRI and fMRI) to obtain the optimal transdiagnostic features. Because the self-reported instruments were administered to acquire a rich set of phenotypic information rather than providing diagnoses, our study also sought to establish the clinical utility of these phenotypic features in distinguishing patients from HCs. In addition, the clinical utility of the markers identified in each feature modality was systematically evaluated by comparing the performance of models trained on each modality. We then report these shared features and discuss the identified latent psychopathological structure across these psychiatric disorders.

## 2 Materials and Methods

### 2.1 The CNP dataset

We utilized the openly available dataset from the CNP LA5c Study conducted at the University of California, Los Angeles (the CNP dataset: https://openneuro.org/datasets/ds000030/versions/00016). Detailed information on the CNP study/dataset can be found in (Poldrack et al., 2016). The CNP dataset contains a variety of data modalities. In this study, we focused on identifying shared transdiagnostic features based on the item-level data from self-reported instruments and neuroimaging data (including both sMRI and resting-state fMRI). The dataset in this study includes 272 subjects, of which 50 are diagnosed with schizophrenia (SCZ), 49 with bipolar disorder (BD), and 43 with attention deficit/hyperactivity disorder (ADHD). The remaining 130 subjects are age-matched healthy controls (HC). The diagnoses were given by following the Diagnostic and Statistical Manual of Mental Disorders, Fourth Edition, Text Revision (DSM-IV-TR; American Psychiatric Association, 2000) and were based on the Structured Clinical Interview for DSM-IV (First et al., 2002). To better characterize ADHD related symptoms, the Adult ADHD Interview (Kaufman et al., 2000) was further used as a supplement. Out of all subjects, 1 had incomplete phenotype data from the instruments used in this study, 10 had missing structural MRI (sMRI) data, and 10 had missing resting-state functional MRI (fMRI) data. Fifty-five (55) subjects had an aliasing artifact in their sMRI data potentially caused by the headset used in the scanner, whereas 22 subjects had errors in the structural-functional alignment step during MRI preprocessing. These subjects were excluded from the corresponding modeling analyses performed in this study. The final number of subjects included in each modeling analysis is as follows: phenotype data, 271; sMRI, 206; fMRI, 229; sMRI+fMRI, 178; phenotype+sMRI, 205; phenotype+fMRI, 228; phenotype+sMRI+fMRI: 177. The demographic information from the subjects are given in **Table 1**.

**Table 1.** Demographic Information*

|  | **HC** | **SCZ** | **BD** | **ADHD** | **Total** |
| --- | --- | --- | --- | --- | --- |
| No. of subjects | 130 | 50 | 49 | 43 | 272 |
| With complete phenotype data | 130 | 50 | 48 | 43 | 271 |
| With sMRI data** | 98 | 30 | 44 | 34 | 206 |
| With fMRI data<sup>†</sup> | 104 | 47 | 41 | 37 | 229 |
| **Age** |  |  |  |  |  |
| Mean age | 31.26 | 36.46 | 35.15 | 33.09 |  |
| SD age | 8.74 | 8.88 | 9.07 | 10.76 |  |
| Range age | 21-50 | 22-49 | 21-50 | 21-50 |  |
| **Gender** |  |  |  |  |  |
| No. of female subjects | 62 | 12 | 21 | 22 |  |
| Percent female subjects | 47.69% | 24.00% | 42.86% | 51.16% |  |
| **Race** |  |  |  |  |  |
| American Indian or Alaskan Native | 19.23% | 22.00% | 6.25% | 0% |  |
| Asian | 1.54% | 2.00% | 0% | 2.33% |  |
| Black/African American | 0.77% | 4.00% | 2.08% | 2.33% |  |
| White | 78.46% | 66.00% | 77.08% | 88.37% |  |
| More than one race | 0% | 2.00% | 14.58% | 6.98% |  |
| **Education** |  |  |  |  |  |
| No high school | 1.54% | 18.00% | 2.08% | 0% |  |
| High school | 12.31% | 44.00% | 29.17% | 23.26% |  |
| Some college | 20.77% | 18.00% | 25.00% | 30.23% |  |
| Associate's degree | 7.69% | 4.00% | 6.25% | 6.98% |  |
| Bachelor's degree | 50.00% | 10.00% | 29.17% | 32.56% |  |
| Graduate degree | 6.92% | 0% | 4.17% | 2.33% |  |
| Other | 0.77% | 4.00% | 4.17% | 4.65% |  |

\* Demographic information is based on initial number of subjects
\*\* Excluding subjects with aliasing artifacts
<sup>†</sup> Excluding subjects with misaligned structural-function imaging data

### 2.2 Phenotype data

It should be noted that we used the term phenotype to broadly refer to behavioral and symptom measures characterized by the clinical instruments. A total of 20 clinical instruments were administered in the CNP dataset to capture a wide range of phenotype data including specific behavioral traits and symptom dimensions (Poldrack et al., 2016). These instruments are either clinician-rated or self-reported. While the clinician-rated questionnaires only covered relevant patient groups, 13 self-reported clinical scales were given to all three patient groups as well as the heathy controls. We therefore selected to only use subjects’ answers to each of the individual questions coming from these 13 self-reported scales as input features to our models because these scales captured phenotypic features across all diagnostic groups as well as the HCs. It should be noted that these 13 self-reported scales were not used to provide the official diagnosis in the CNP data since they are not designed for such purposes. Specifically, the 13 self-reported scales used in this study are: Chapman Social Anhedonia Scale, Chapman Physical Anhedonia Scale, Chapman Perceptual Aberrations Scale, Hypomanic Personality Scale, Hopkins Symptom Checklist, Temperament and Character Inventory, Adult ADHD Self-Report Scale v1.1 Screener, Barratt Impulsiveness Scale, Dickman Functional and Dysfunctional Impulsivity Scale, Multidimensional Personality Questionnaire – Control Subscale, Eysenck’s Impulsivity Inventory, Scale for Traits that Increase Risk for Bipolar II Disorder, and Golden and Meehl’s Seven MMPI Items Selected by Taxonomic Method. Together, these self-reported scales cover domains including symptoms, personality traits, positive and negative affect, cognition, as well as sensory and social processing. The scores for known sum and sub-scores within these self-reported instruments for each patient group as well as the HCs are reported in **Supplementary Table 1**.

### 2.3 MRI data acquisition parameters

MRI data were acquired on one of two 3T Siemens Trio scanners both housed at the University of California, Los Angeles. The sMRI data used in this study are T1-weighted and were acquired using a magnetization-prepared rapid gradient-echo (MPRAGE) sequence with the following acquisition parameters: TR = 1.9 s, TE = 2.26 ms, FOV = 250 mm, matrix = 256 × 256, 176 1-mm thick slices oriented along the sagittal plane. The resting-state fMRI data contain a single run lasting 304 s. The scan was acquired using a T2*-weighted echoplanar imaging (EPI) sequence using the following parameters: 34 oblique slices, slice thickness = 4 mm, TR = 2 s, TE = 30 ms, flip angle = 90°, matrix size 64 × 64, FOV = 192 mm. During the resting-state scan, subjects remained still and relaxed inside the scanner, and kept their eyes open. No specific stimulus or task was presented to them.

### 2.4 MRI preprocessing

#### 2.4.1 sMRI

Structural MRI preprocessing was implemented using Freesurfer’s *recon-all* processing pipeline (http://surfer.nmr.mgh.harvard.edu/). Briefly, the T1-weighted structural image from each subject was intensity normalized and skull-stripped. The subcortical structures, white matter, and ventricles were segmented and labeled according to the algorithm described in (Fischl et al., 2002). The pial and white matter surfaces were then extracted and tessellated (Fischl et al., 2001), and cortical parcellation was obtained on the surfaces according to a gyral-based anatomical atlas which partitions each hemisphere into 34 regions (Desikan et al., 2006).

#### 2.4.2 Resting-state fMRI

Resting-state fMRI preprocessing was implemented in AFNI (http://afni.nimh.nih.gov/afni). Specifically, the first 3 volumes in the data were discarded to remove any transient magnetization effects in the data. Spikes in the resting-state fMRI data were then removed and all volumes were spatially registered with the 4^th^ volume to correct for any head motion. The T1w structural image was deobliqued and uniformized to remove shading artifacts before skull-stripping. The skull-stripped structural image was then spatially registered with motion corrected fMRI data. The fMRI data were further spatially smoothed using a 6-mm FWHM Gaussian kernel and converted to percent signal change. Separately, the Freesurfer-generated aparc+aseg image from sMRI preprocessing was also spatially registered with and resampled to have the same spatial resolution of the BOLD image. Based on this, eroded white matter and ventricle masks were created, from which nuisance tissue regressors were built based on non-spatially smoothed fMRI data to model and remove variances that are not part of the BOLD signal. Specifically, we used the ANATICOR procedure (Jo et al., 2010), where a locally averaged signal from the eroded white matter mask within a 25-mm radius spherical region of interest (ROI) centered at each gray matter voxel was used to create a voxel-wise local estimate of the white matter nuisance signal. This local estimate of the white matter nuisance signal, along with the estimated head motions and average signal from the ventricles were detrended with a 4^th^ order polynomial and then regressed out from the fMRI data. Finally, the clean resting-state fMRI data was spatially normalized to the MNI template and resampled to have 2 mm isotropic voxels. Note that in our preprocessing pipeline, the spatial normalization was performed after regressing out nuisance signals. This allowed nuisance tissue regression to be performed in each subject’s native space to achieve a more accurate removal of these signals.

### 2.5 Feature extraction

We extracted measures from 3 data modalities as features: phenotype data from self-reported instruments, measures derived from the sMRI data, and functional correlations based on resting-state fMRI data. For phenotype features from self-reported instruments, we directly used subjects’ responses from a total of 578 questions from the above listed 13 instruments. Responses from non-True/False type questions were normalized to have a range of between 0 and 1 to match those from True/False type questions. For sMRI features, we specifically used 1) the volume of subcortical structures generated by Freesurfer’s subcortical volumetric segmentation, and 2) the area, thickness, and volume of cortical brain regions estimated from Freesurfer’s surface-based analysis pipeline. For resting-state fMRI features, we first parceled the brain into 264 regions according to the atlas proposed in (Power et al., 2011). A 5-mm radius spherical ROI was seeded according to the MNI coordinates of each brain region specified in the atlas. Second, the clean resting-state BOLD time series from all voxels within a given 5-mm radius spherical ROI were averaged to create the representative time series for the brain region. Third, functional connectivity between ROIs was estimated via the Pearson’s correlation coefficient between the average time series from all pairs of brain regions. This produced a 264-by-264 correlation matrix, from which 34,716 are unique correlations between two distinct ROIs and were used as input features to the models.

### 2.6 Model fitting and feature importance weighting

The primary goals of machine learning analyses in this study were two-fold: 1) to identify important features shared across a transdiagnostic patient group and 2) to evaluate the clinical utility of the transdiagnostic features via classifiers that can reliably separate the patient group as a whole from healthy controls. To achieve these goals, we built classifiers based on the logistic regression model as implemented in the *scikit-learn* toolbox to classify patients from HCs. To identify predictive transdiagnostic features embedded within each feature modality, separate logistic regression models were independently trained using each of the above extracted feature modalities (i.e., item-level phenotype data, sMRI measures, and resting-state fMRI correlations) as inputs and their performances were evaluated in each of the transdiagnostic scenarios. Combinations of 2 and 3 feature modalities were also used as classifiers’ inputs and their performances were evaluated in the same fashion.

Because the number of features we extracted was relatively large compared to the sample size in CNP data, an elastic net regularization term (Zou and Hastie, 2005) was added in all of our logistic regression models to prevent overfitting. The use of elastic net regularization in our models also enabled feature selection as the regularization induces sparse models via the grouping effect where all the important features will be retained and the unimportant ones set to zero (Zou and Hastie, 2005; Ryali et al., 2012). This allowed us to identify predictive features that are shared across multiple patient categories.

We adopted the following procedure to determine the best regularization parameters. First, the input data were randomly partitioned into a development set and an evaluation set. The development set contains 80% of the data upon which a grid search with 3-fold cross validation procedure was implemented to determine the best regularization parameters. Then the model with the best regularization parameters was further tested on the remaining 20% of evaluation set. All features were standardized to have zero mean and unit variance within the training data and the mean and variance from the training data were used to standardize the corresponding test data. The entire process was implemented 100 times. The following metrics were used to quantify the model performances: area under the receiver operating characteristics curve (AUC), accuracy, sensitivity, and specificity. The mean and standard deviation of the above metrics over the 100 evaluation sets were reported.

From the above models, the predictive power of each feature is assessed via the weights of the logistic regression model in our transdiagnostic classifiers. For each feature, we calculated its corresponding standardized model weight (mean model weight divided by the standard deviation) across the 100 model implementations as the proxy for feature importance. Features with large importance values from our transdiagnostic classifiers are potentially symptoms, traits, and neuropathological mechanisms shared across patient groups but are distinct from healthy controls.

To identify the set of most predictive transdiagnostic features within a given data modality, we used the following feature importance-guided sequential model selection procedure. Specifically, we first rank ordered the features in the classifiers according to their standardized model weights. Next, a series of truncated models was built such that each model only takes the top k most predictive features as inputs to perform the same classification tasks. The model weights and the best regularization parameters for each truncated model were estimated via the 3-fold cross validation procedure within the development set. We let k range from the top 1 most predictive feature to all available features in steps of 1 for phenotype features, sMRI features, and the combination of the two feature sets. For any feature or feature combinations involving fMRI correlations, because of the significantly increased feature dimension, the k’s were chosen from a geometric sequence with a common ratio of 2 (i.e., 1, 2, 4, 8, 16, …). The sequential model selection procedure was implemented 10 times for each feature set. Model performances based on the evaluation set were obtained for each truncated model and were evaluated as a function of the number of top features (k) included in each truncated model to determine the optimal feature set.

### 2.7 Statistical analyses

To statistically examine whether the models’ performances are significantly above chance level, we performed a random permutation test where labels in the training data (e.g., HC vs. Patients) were shuffled 100 times and truncated models based on the best set of features were trained on these label-shuffled data using exactly the same approach as described above (Ojala and Garriga, 2009). The performances from the 100 models were used to construct the empirical null distribution against which the performance of the best truncated models based on the actual unshuffled data was then compared. This random permutation test procedure also helped us to determine whether overfitting occurred during training.

To evaluate differences in sum scores obtained from the top features between HC and patients, we used two sample t-tests on the sample means since the sum scores are quasi normally distributed. Effect sizes were measured using Cohen’s d, which captures the shift in mean scaled by the data’s standard deviation. Tests on the difference in AUC between the full model and the best truncated model were carried out via the Wilcoxon’s rank-sum test.

## 3 Results

In total, we trained classifiers to distinguish HCs from the patient group as a whole based on 7 sets of features by either using each individual feature modality (self-reported instruments, sMRI, and fMRI) or combinations of 2 or 3 feature modalities (e.g., instruments+sMRI+fMRI). The classifiers’ performances using each of the 7 feature sets for the HC vs. Patients classifier are reported in **Table 2**. Overall, classifiers trained on feature sets involving phenotypical data from self-reported instruments (i.e., scales and scales + MRI feature sets) outperformed those only trained on MRI features (sMRI, fMRI, and sMRI+fMRI). For classifiers using features involving these instruments, the mean AUC ranged from 0.83 to 0.89 (mean accuracy: 0.77 – 0.91), whereas the mean AUC ranged from 0.56 to 0.59 (mean accuracy: 0.58 – 0.61) for MRI feature sets.

**Table 2.** Performances of the transdiagnostic models on each feature set*

| **Performance of the full model:** |  |  |  |  |  |  |  |
| --- | --- | --- | --- | --- | --- | --- | --- |
|  | Scales | sMRI | fMRI | s+fMRI | Scales+sMRI | Scales+fMRI | Scales+s+fMRI |
| AUC | 0.83(0.04) | 0.56(0.05) | 0.59(0.04) | 0.57(0.05) | 0.89(0.07) | 0.86(0.06) | 0.86(0.05) |
| Accuracy | 0.77(0.05) | 0.58(0.08) | 0.60(0.06) | 0.61(0.07) | 0.91(0.04) | 0.87(0.05) | 0.86(0.05) |
| Sensitivity | 0.77(0.07) | 0.74(0.11) | 0.60(0.11) | 0.62(0.11) | 0.86(0.08) | 0.82(0.12) | 0.86(0.08) |
| Specificity | 0.82(0.07) | 0.38(0.11) | 0.56(0.12) | 0.49(0.17) | 0.80(0.15) | 0.61(0.30) | 0.48(0.26) |
| **Performance of the best truncated model:** |  |  |  |  |  |  |  |
|  | Scales | sMRI | fMRI | s+fMRI | Scales+sMRI | Scales+fMRI | Scales+s+fMRI |
| AUC | 0.95(0.02) | 0.78(0.06) | 0.87(0.08) | 0.77(0.06) | 0.96(0.03) | 0.98(0.02) | 0.96(0.03) |
| Accuracy | 0.88(0.04) | 0.71(0.06) | 0.85(0.07) | 0.77(0.06) | 0.87(0.05) | 0.92(0.04) | 0.90(0.04) |
| Sensitivity | 0.87(0.08) | 0.81(0.09) | 0.86(0.09) | 0.77(0.08) | 0.93(0.07) | 0.91(0.06) | 0.94(0.05) |
| Specificity | 0.88(0.04) | 0.60(0.16) | 0.84(0.18) | 0.76(0.07) | 0.80(0.15) | 0.92(0.04) | 0.85(0.09) |
| No. of features | 85 | 131 | 8192 | 16384 | 238 | 32 | 64 |
| **Test on AUCs between the full and the truncated model\*\*:** |  |  |  |  |  |  |  |
|  | Scales | sMRI | fMRI | s+fMRI | Scales+sMRI | Scales+fMRI | Scales+s+fMRI |
| Test statistic | 100 | 100 | 100 | 99.5 | 82.5 | 100 | 95 |
| p-value<sup>†</sup> | < 0.001 | < 0.001 | < 0.001 | < 0.001 | 0.01537 | < 0.001 | < 0.001 |

\* The mean performance measures across 10 implementations are reported here with the standard deviation shown in parentheses
\*\* Wilcoxon's rank-sum test
† FDR-corrected

Next, to identify the optimal set of transdiagnostic features shared across the CNP patient population that are highly distinct from HC, we examined the performance measures from the best truncated classification models during sequential model selection (**Figure 1** and **Table 2;** see **Supplementary Fig. 1** for AUC as a function of input feature dimensions). Significantly improved performances were obtained from the best truncated classification models compared with the corresponding models using the full sets of features (all p’s < 0.05, FDR corrected, as assessed by the rank-sum test; **Table 2**). The AUCs from all feature sets were also significantly above chance level as assessed via the random permutation test (all p’s < 0.01, FDR corrected; **Supplementary Fig. 2**). Additionally, the computational time for the importance guided sequential model selection method grew linearly as the number of features increased, which is highly efficient compared to the brute force feature selection procedure (exponential time complexity; **Supplementary Fig. 3**).

The truncated classification model involving data from the self-reported instruments alone had high performance of distinguishing patients from HCs with the mean AUC being 0.95 (accuracy: 0.88; sensitivity: 0.87; specificity: 0.88; **Figure 1**; **Table 2**). This truncated model selected 85 items as the most predictive features from the total of 578 items contained in the 13 self-reported instruments. Moreover, only 10 items were needed to achieve an AUC of 0.90 (accuracy: 0.81; sensitivity: 0.79; specificity: 0.84), suggesting that a concise scale can be constructed potentially for screening purposes. The model involving data from self-reported instruments performed better compared to those using feature sets based solely on MRI (mean AUC ranging from 0.77 to 0.87; mean accuracy ranging from 0.71 to 0.85). Combining MRI features with data from instruments only slightly improved the model performance (mean AUC being 0.96 – 0.98) (**Figure 1**; **Table 2**). Taken together, this indicates that the phenotypical data captured by the 13 self-reported instruments contain a set of transdiagnostic features common across the patient populations studied while distinguishing them from the healthy controls. We hence focused on discussing these transdiagnostic phenotypical features below.

**Figure 1.**
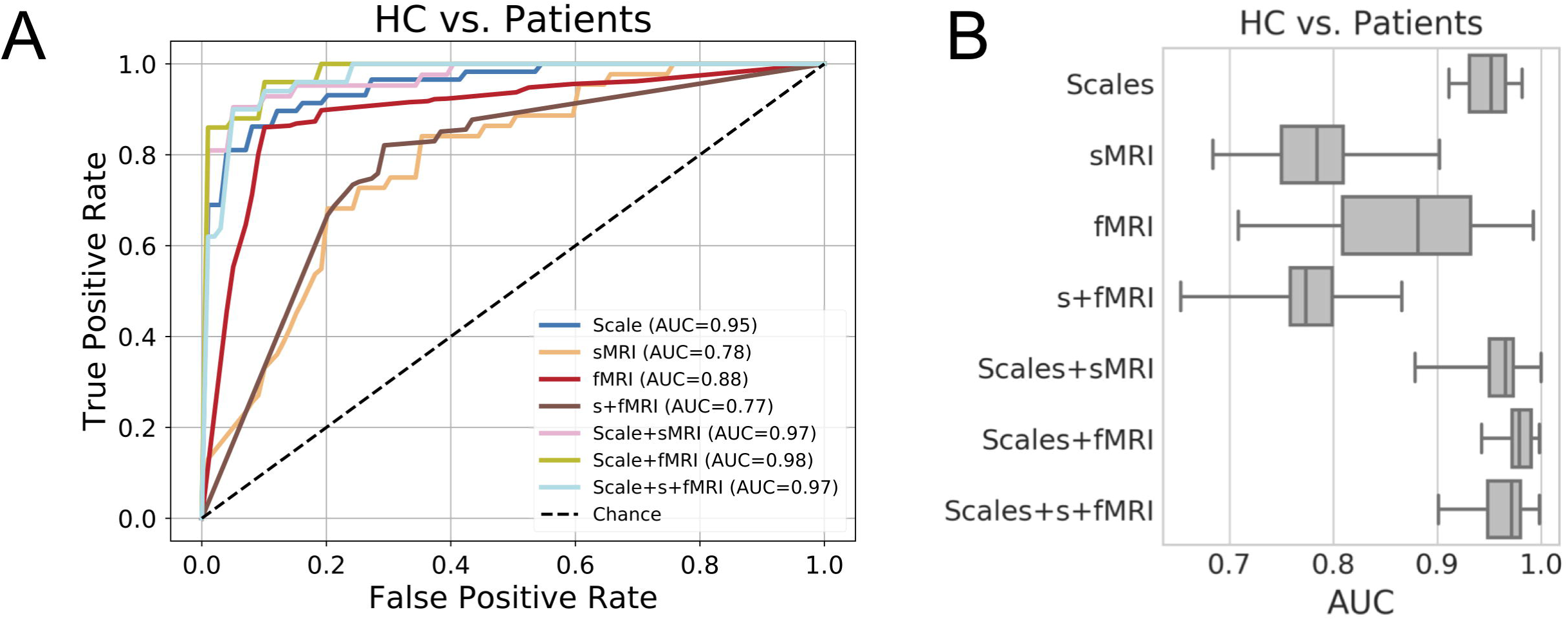
The performances of the best transdiagnostic models selected via the feature importance-guided sequential model selection procedure. **A)** The receiver operating characteristic (ROC) curve for the best truncated models based on each feature set. Area under the ROC curve (AUC) for each model is listed in the legend. **B)** Box plots showing the AUC of the best truncated model for each feature set measured across 10 implementations of sequential model selection procedure.

**Figure 2.**
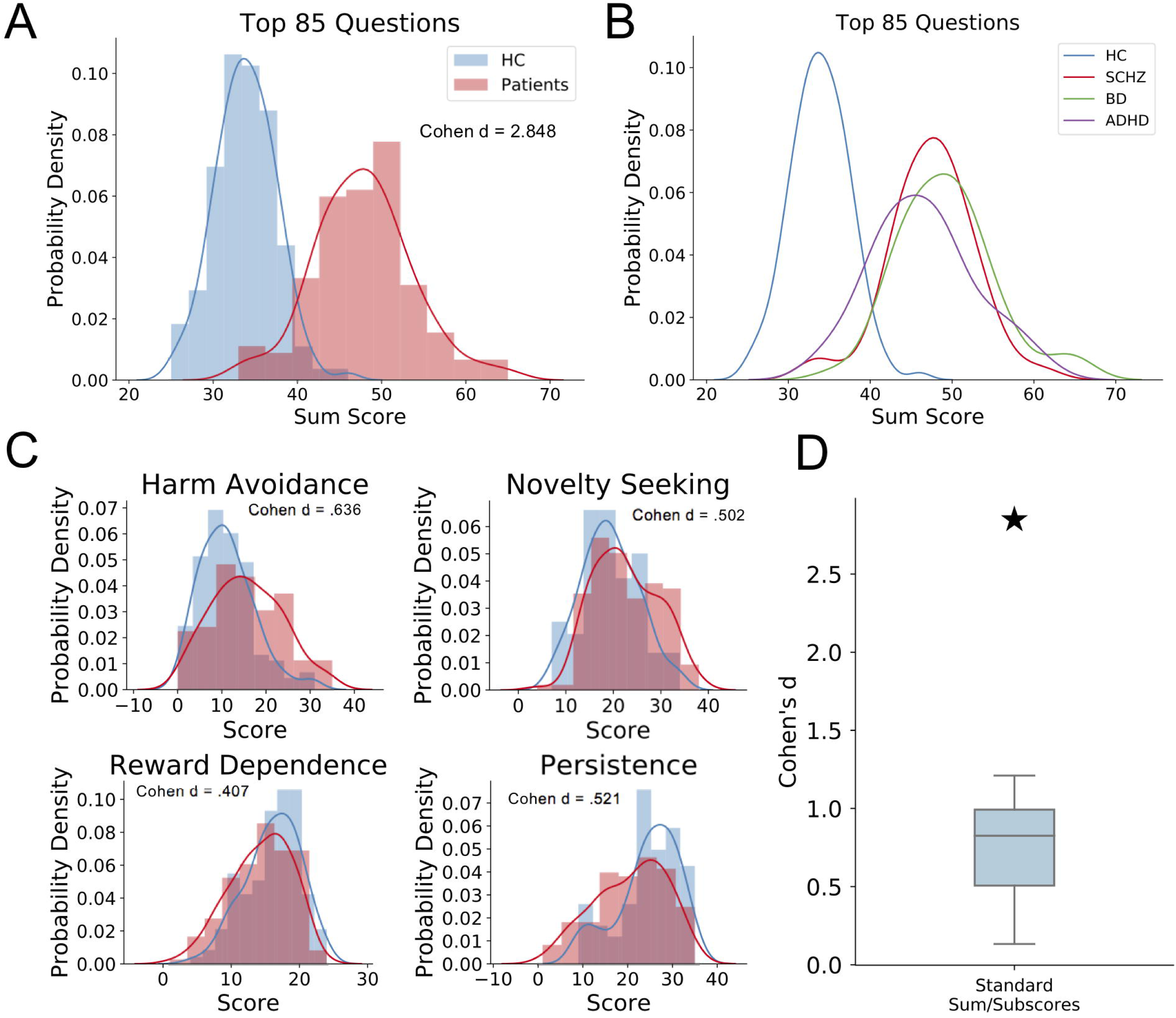
Distributions and effect sizes of the model’s derived scores vs. the existing scale scores. **A)** Sum score calculated from the identified 85 most predictive items showing high separability in terms of Cohen’s d between HC and Patients. **B)** All three patient categories showed elevated sum scores relative to HC (p < 0.001). **C)** The 4 temperament sub-scores in TCI included in the CNP dataset showing only medium effect sizes between HC and Patients. **D)** Box plot showing significantly higher effect size from the identified 85 items (asterisk) compared to all predefined sum and subscores in self-reported instruments in CNP data. The asterisk represents the Cohen’s d between HC and Patients from the top 85 items, whereas the box plot shows the effect sizes from all predefined sum and subscores (also see Supplementary Table 2).

**Figure 3.**
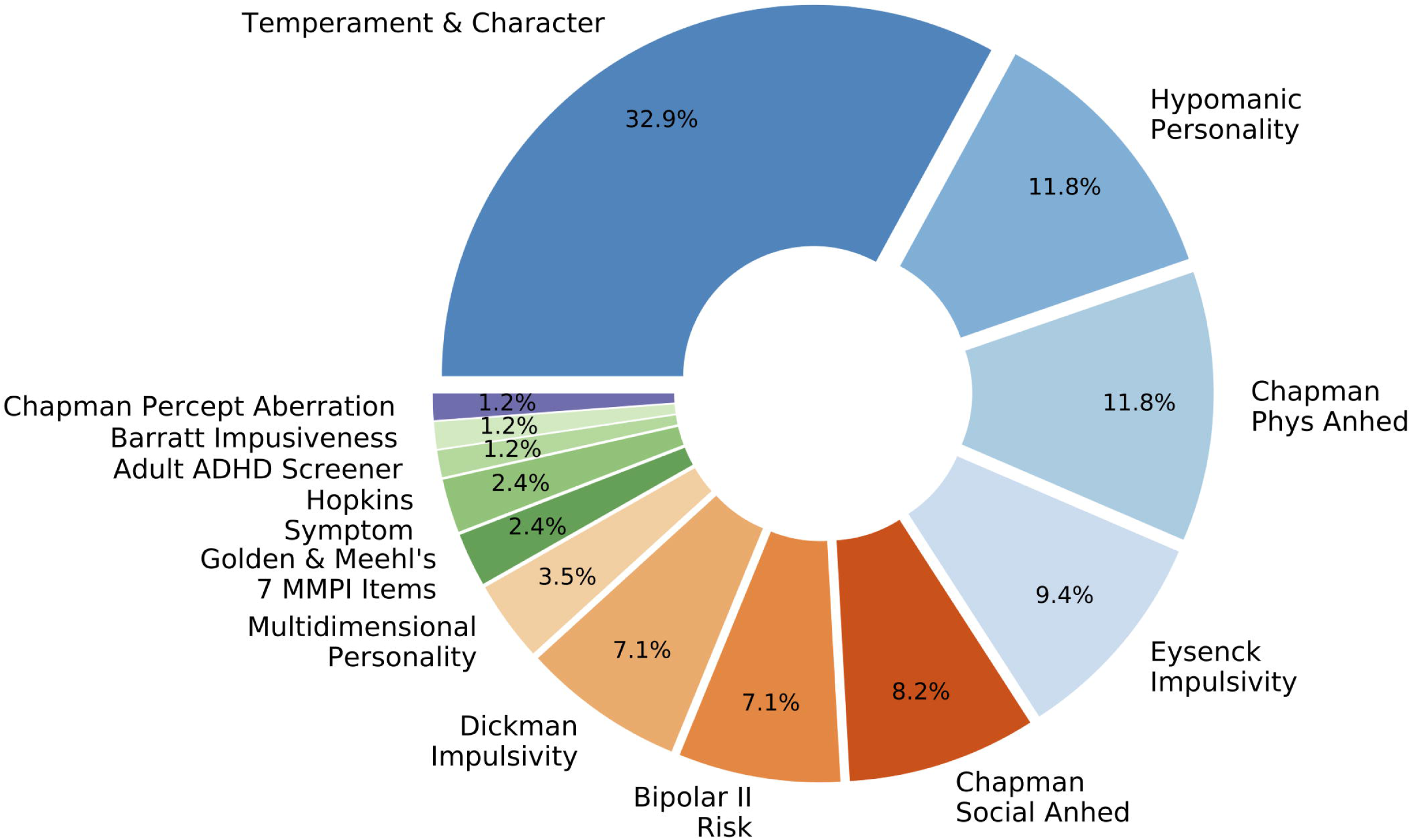
The percentage of items from each of the 13 self-reported instruments among the set of 85 most predictive transdiagnostic items.

It should be noted that small yet systematic differences exist between patients and HCs in the CNP dataset despite the effort to fully counterbalance subjects’ demographic information during recruiting. From the demographic information listed in **Table 1**, a statistical test on age showed that the median age for patients were significantly higher than HCs (Mann-Whitney’s U test: p = 0.001). The gender ratio on the other hand did not significantly differ between patients and HCs (Z-test on proportions: p = 0.14). Patients had significantly lower years of education compared with HCs (Mann-Whitney’s U test: p < 0.001). The racial distribution also showed a significant difference between patients and HCs (*χ*^2^-test: p = 0.004). To ensure that the results were not mainly driven by differences in demographics, we built classifiers based solely on these demographic variables. The mean test AUC on the evaluation set for the best model was 0.71 (SD = 0.06), which was substantially lower than the performance of both the full model and the best truncated model based on the self-reported instruments alone (mean AUCs being 0.83 and 0.95, respectively). Additionally, despite mixing the demographic variables with the self-reported instrument items slightly improved model performance compared to those obtained from instruments alone (mean AUC for truncated model using demographics and instruments: 0.98), the highest-ranking demographic variable ranked only 43^rd^ among all instrument items and demographic features. Taken together, these results suggest that our models were unlikely to be mainly driven by demographic differences.

A simple sum score constructed by adding up an individual’s responses to the 85 most predictive items (with item responses having negative model coefficients reversed) demonstrated high separability between healthy controls and patients (Cohen’s d = 2.85; test on the difference in sample mean: t = 10.27, p < 0.001; **Figure 2A**). The separability based on the top 10 most predictive items (corresponding AUC = 0.90) was also very high (Cohen’s d = 2.16; test on the difference in sample mean: t = 18.13, p < 0.001). The sum scores had higher separability than other known sum scores/sub-scores of the self-reported instruments (**Figure 2C & 2D; Supplementary Table 2**), indicating the items selected by the sequential model selection procedure do not adhere to known dimensional structures within the instruments. Calculating the sum score for each individual patient category showed that all 3 patient categories had elevated sum scores compared to healthy controls (t 5.671, p < 0.001); yet, the difference between the patient categories was insignificant (t < 1.940, p 0.056; **Figure 2B**). This suggests that the 85 items captured transdiagnostic phenotypic features shared across the patient groups as a whole rather than driven by a single patient category.

**Figure 3** illustrates the proportion of questionnaire items selected from each instrument that were included in transdiagnostic set of phenotypic features of the best truncated model (i.e. the one with the highest AUC). Items from all 13 instruments were selected to be among the top features by the classifiers. Overall, these instruments measure a wide range of phenotype and symptom domains encompassing personality traits, positive and negative affect (reward/anhedonia, fear, and anxiety), cognition (attention, response inhibition), sensory processing (perceptual disturbances), and social processing. While all items included among the set of transdiagnostic phenotypic features jointly formed a highly predictive set to distinguish patients from healthy controls, the Temperament and Character Inventory (TCI) contributed the largest proportion of items in the set of transdiagnostic features. The proportion of TCI items selected among the 85 most predictive items (32.9%) significantly exceeded the proportion of all TCI items among all 578 items from the 13 instruments (24.0%; p = 0.04 as assessed via the Z-test). The disproportionately high number of TCI items indicates that certain personality traits are strong predictors of shared psychopathology regarding SCZ, BD, and ADHD.

To better understand which features strongly predicted psychopathology, we focused on the top 20 items that contributed the largest magnitude of model weights. We identified the specific behavioral/symptom phenotypes characterized by each item and then grouped the items accordingly. Among personality traits, the top transdiagnostic phenotypes included neuroticism, extraversion, and impulsivity (**Figure 4A**); whereas the top symptom domains consisted of mood dysregulation, inattention, hyperactivity/agitation, and social anhedonia and apathy. In addition, the importance of religion was also a shared feature across patients (**Figure 4A**; see **Supplementary Table 4** for item grouping). We next compared the most predictive transdiagnostic features with those most predictive of a single patient category from healthy controls to identify category-specific differences (**Figure 4B-D**; see **Supplementary Table 3 and Supplementary Fig. 4** for classification results between HC and each patient category). SCZ patients exhibited additional features including perceptual aberration, physical anhedonia, and psychological distress that are not among the top transdiagnostic features. On the other hand, extraversion, impulsivity, inattention, and religion which were present in the transdiagnostic features set were not among the most predictive features for SCZ (**Figure 4B** and **Supplementary Table 5**). By contrast, transdiagnostic features overlapped with BD patient-specific features. Nonetheless, BD patients exhibited additional features of increased energy, psychological distress and physical anhedonia; yet psychomotor agitation and neuroticism contributed little predictive value (**Figure 4C**, **Supplementary Table 6**). ADHD patients were effectively classified by additional features such as indecision and physical anhedonia, with little predictive contribution from apathy, neuroticism, and religion (**Figure 4D**, **Supplementary Table 7**).

**Figure 4.**
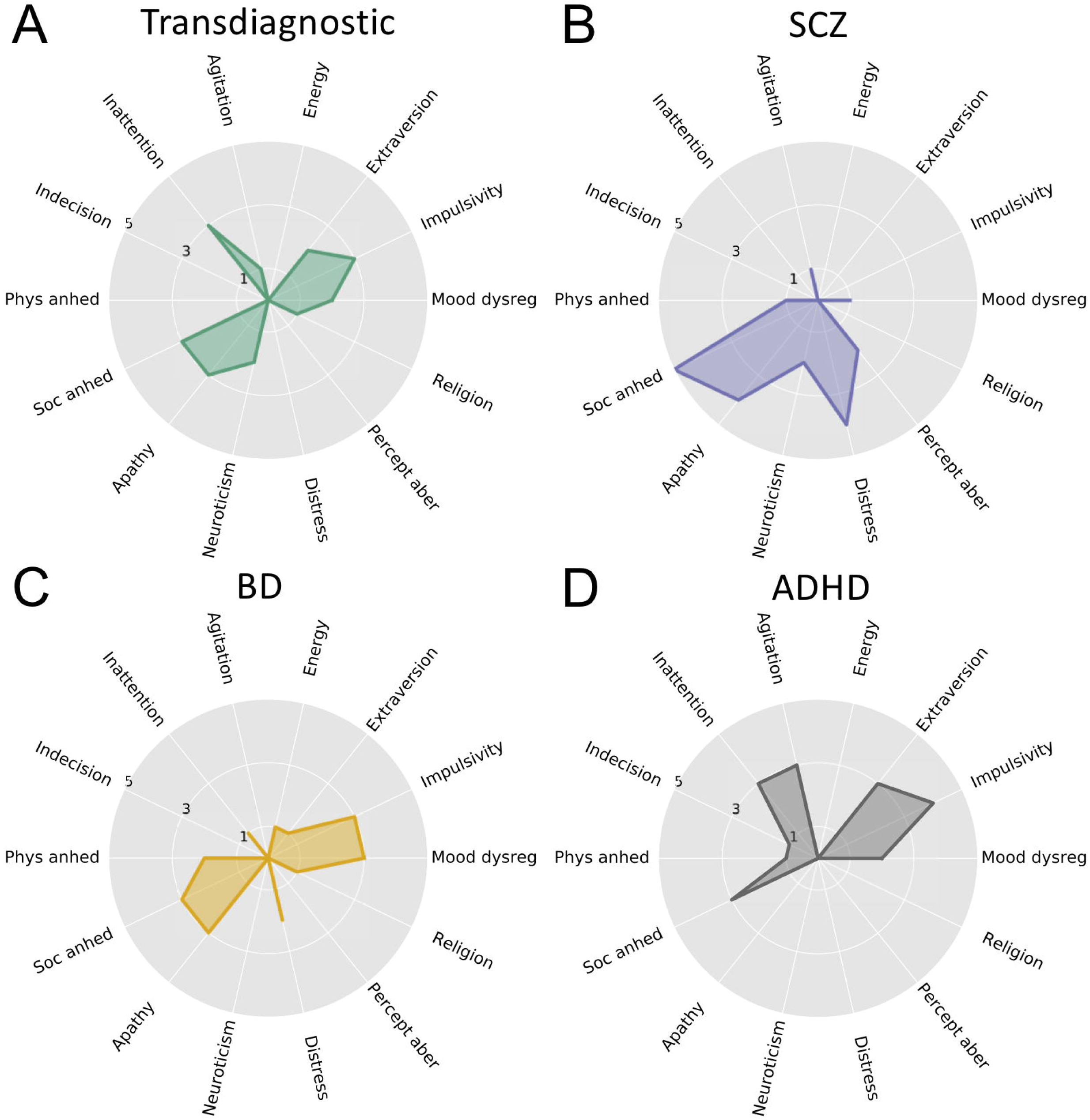
The grouping of items into specific phenotypic domains for the top 20 most predictive items from **A)** the HC vs. All Patients transdiagnostic model and **B) – D)** the 3 HC vs. a single patient category classifiers. The radius in the spider plots represents item counts.

## 4 Discussion

In this study, using self-reported instruments provided in the CNP dataset, we generated predictive models to identify a set of transdiagnostic phenotypic features that were shared across SCZ, BD, and ADHD. These models were quantified for performance (e.g. accuracy, sensitivity and specificity) and were interpretable along dimensions of personality traits and symptom domains. We found the set of 85 items is highly predictive of the patient group as a whole from HCs. To our surprise, a compact model of only 10 items is sufficient to achieve a performance AUC value of 0.90. Further, we demonstrated that a simple sum score can be calculated to enable high separability between patients and HCs. Our importance-guided sequential model selection approach revealed which phenotypical features were shared across transdiagnostic patient groups. Within each patient population, we also show which abnormal psychopathological personality traits and symptom domains deviated from the transdiagnostic classifier. Importantly, many of these features are consistent with established clinical intuition. Taken together, this study offers new perspectives on the shared psychopathology across SCZ, BD, and ADHD and underscores the potential of creating a short transdiagnostic screening scale based on the selected items.

The application of machine learning to systematically search for consistent patterns in clinical data across disease categories defined in DSM is an emerging trend in the field of computational psychiatry (Bzdok and Meyer-Lindenberg, 2017). Nonetheless, our approach to identifying transdiagnostic features in psychiatric disorders differs both conceptually and methodologically from previous studies. Numerous investigators have focused on patient subtyping within a given disorder (Rhebergen et al., 2011; Lamers et al., 2012; van Loo et al., 2012, 2014; Georgiades et al., 2013; Brodersen et al., 2014; Doshi-Velez et al., 2014; Lewandowski et al., 2014; van Hulst et al., 2014; Veatch et al., 2014; Costa Dias et al., 2015; Geisler et al., 2015; Sun et al., 2015; Clementz et al., 2016; Drysdale et al., 2016; Mostert et al., 2018) or have mined transdiagnostic symptom dimensions underlying various psychiatric disorders (Grisanzio et al., 2017; Elliott et al., 2018; Xia et al., 2018a, b). Among studies examining the transdiagnostic symptom dimensions, most adopted an unsupervised machine learning predictive framework. However, the differences in distance/similarity metrics used, coupled with the lack of ground truth in the unsupervised machine learning algorithms used to detect the transdiagnostic structure, make it difficult to validate the clinical utility of the identified features. We designed our study to overcome these limitations. To our best knowledge, our study is the first to use feature importance to guide forward model selection under a supervised machine learning framework to identify transdiagnostic psychopathological features across multiple DSM categories. The high performance of our truncated models selected via the model selection approach demonstrate the potential clinical utility of the identified transdiagnostic features.

Though we built models with different modalities as inputs (e.g. personality traits, symptoms and neuroimaging), we found high performance models could be obtained without significant contribution of the imaging modalities. This finding contrasts with what would be predicted from the published literature. For example, a recent meta-analysis of studies on psychiatric disorders involving structural magnetic resonance imaging (sMRI) identified shared abnormalities in certain brain regions underlying common psychiatric disorders (Goodkind et al., 2015). In addition, studies using functional MRI (fMRI) found altered functional connectivity patterns shared across multiple categories of disorders such as SCZ, BD, and major depressive disorder (MDD) (Buckholtz and Meyer-Lindenberg, 2012; Wei et al., 2018). Similarly, another recent study focusing on MDD, post-traumatic stress disorder, and panic disorder identified 6 distinct subtypes based on 3 orthogonal symptom dimensions shared across the DSM diagnoses and their corresponding biomarkers in electroencephalogram (EEG) beta activity (Grisanzio et al., 2017). Although these studies did not systematically compare the predictability in each data modality, it is possible that the sample size in CNP or other methodological differences (e.g., parcellation used during sMRI and fMRI feature extraction) limited the weighted importance of structural or functional measures in our models.

A broad set of behavioral phenotypic features from the self-report instruments were identified by our sequential model selection procedure to be shared across the three patient groups. The phenotypes are distributed across all 13 self-reported instruments and covers symptom domains encompassing personality and traits, positive and negative affect, cognition, sensory and social processing. It should be noted that these 13 self-reported instruments are not designed to yield diagnoses. Therefore, a set of phenotypic features that can be identified algorithmically and used to distinguish patients from HCs with an accuracy level close to a clinician’s performance demonstrates the potential clinical utility of the phenotypic features and the sequential model selection approach. For the top 20 most predictive features, mood dysregulation, impulsivity, inattention, neuroticism, social anhedonia and apathy weighted prominently in the transdiagnostic model. This high level of shared symptom domains across SCZ, BD, and ADHD is in line with recent genetic studies reporting significantly correlated risk factors for heritability among these three disorders (Larsson et al., 2013; The Brainstorm Consortium et al., 2017; Bipolar Disorder and Schizophrenia Working Group of the Psychiatric Genomics Consortium et al., 2018). For SCZ and BD, previous studies have identified shared features both in terms of symptoms and the underlying psychopathology and biology (Pearlson, 2015). Similarly, studies have identified shared symptoms and biology between SCZ and ADHD (Peralta et al., 2011; Park et al., 2018) and have found high levels of comorbidity between BD and ADHD along with the shared features between the two disorders (Nierenberg et al., 2005; Klassen et al., 2010; van Hulzen et al., 2017; Wang et al., 2017). Despite these prior studies, the three diagnostic categories have not been considered together in a single study. Consistent with the findings reported in these studies, our study provides an important data-driven confirmation on the shared behavioral and symptoms features across the three disease categories.

Since the TCI is less commonly used in clinical practice and historically a greater emphasis has been placed on symptoms than personality traits, we were surprised by the finding that the TCI contributed the largest proportion of questions among the set of 85 most predictive items determined by the transdiagnostic classifier. Prior studies have found that the personality traits and characters defined in the TCI are associated with various mood disorders (Cloninger et al., 1998; Grucza et al., 2003). Specifically, for disorders in the CNP dataset, studies have found positive association between personality dimensions characterized in TCI and overall ADHD symptom (Lynn et al., 2005; Anckarsäter et al., 2006) as well as subtypes of ADHD (Salgado et al., 2009). For SCZ, studies have identified links between positive and negative symptom dimensions and TCI factors (Guillem et al., 2002; Hori et al., 2008). Among BD patients, (Hajirezaei et al., 2017) identified personality profiles that are distinct from healthy controls and these profiles were further found to be shared with MDD. Since these studies associated disease symptoms solely with the known factor scores in the TCI, the contribution of the nuanced personality profiles captured in individual items in the TCI could not be determined. In the current study, the fact that we identified items in the TCI that corresponded to shared symptoms such as apathy, anhedonia, and distress extends prior literature and is consistent with studies documenting the relationship between TCI factor scores and symptoms such as anhedonia (Martinotti et al., 2008), as well as depression and anxiety (Jylhä and Isometsä, 2006). Additionally, prior studies only examined TCI’s association with symptoms without simultaneously including other instruments as covariates in the model. Such an approach cannot evaluate the relative importance of the personality traits in TCI against the broader set of phenotypical features defined in other instruments. In this regard, our study established the usefulness of personality traits as a set of reliable transdiagnostic features among all features defined in the self-reported instruments in the CNP data.

The sum score of the 85 most predictive transdiagnostic items achieved much higher separability between HC and patients than known sub-scores and sum scores in the instruments that were specifically designed to assess diagnosis-specific symptom domains. This is true even for the subset of top 10 most predictive transdiagnostic items, which indicates that the shared phenotypic features across patient groups do not fully adhere to known dimensional structures in the instruments. Thus, using the total score and/or the sub-scores according to pre-defined subscales of a given instrument cannot identify the optimal set of transdiagnostic features. One explanation for this phenomenon is that because the patients share a broad range of phenotypic features, the pre-defined subscales and sum scores become insufficient in capturing the full dimensional structure since most of the instruments are designed to measure a limited set of constructs targeting a specific patient population (Avila et al., 2015). This further demonstrates the advantage of our importance-guided sequential model selection approach in identifying potentially clinically relevant transdiagnostic features across a large set of instruments.

While patients shared a broad set of phenotypic features, our results showed deviations from this transdiagnostic structure within the most predictive features for each patient group. These differences may in particular reflect clustered personality traits and symptom domains that are most unique for each patient population. For SCZ, the unique features of perceptual aberration and psychological distress, along with other features that are consistent with the transdiagnostic structure, largely conform to the positive and negative symptom dimensions associative with SCZ patients. For BD, the unique features of increased energy, physical anhedonia, and psychological distress serve to shape the symptom structure along the manic and depressive dimensions. For ADHD, the increased representation in inattention, hyperactivity, and impulsivity is consistent with the overall symptomatology of ADHD patients. Overall, the concurrent existence of shared and category-specific phenotypic features across the CNP patient groups is consistent with recent studies reporting both shared and distinct properties in functional brain networks (Grisanzio et al., 2017; Xia et al., 2018a, b) and genetic neuropathology (The Brainstorm Consortium et al., 2017; Gandal et al., 2018) across major psychiatric disorders. Our results raise the possibility of exploring the relationship between the predictive phenotypic features and the underlying genetics of the individuals or groups that present with these features.

In conclusion, we identified a set of transdiagnostic phenotypic features shared across SCZ, BD, and ADHD. This set of features distinguished the patient group from HC with high accuracy and a compact transdiagnostic screening scale can be derived from the corresponding top 10 most predictive questionnaire items. The feature importance guided sequential model selection provides a data-driven method to identify shared features under a supervised machine learning framework, in which the performance of the identified feature sets is evaluated on unseen data. This is an advantage over unsupervised machine learning methods. Moreover, the importance guided sequential model selection can be generalized to identify clinically-useful transdiagnostic features across categories defined in DSM-5 and ICD-10, or alternatively to identify the neural correlates of symptom severity across psychiatric disorders (Mellem et al., 2018). It should be noted that the medication status in the CNP dataset is not controlled. This suggests that although reliable transdiagnostic features could be identified across patient groups, the underlying cause of the observed symptom structure could potentially be confounded by the uncontrolled medication and symptom status. Future studies should further validate the transdiagnostic features identified in this study on other datasets with similar patient populations and with better controlled medication status. Including these additional datasets as out-of-sample validations can demonstrate the generalizability of the current results and methodology to the wider population. Additionally, because the phenotypic features largely reflected behaviors and symptoms through self-reports, the high performance of our models based on these features may be reflecting the close mapping between the identified features and symptom-based diagnostic criteria in DSM. Nevertheless, our findings represent an important step in the ongoing effort to characterize clinically useful transdiagnostic phenotypes. Future studies could potentially investigate the underlying brain circuits associated with these clinically-relevant phenotypic features.

## Supporting information

## 6 Author Contributions

YL, MM, HG, WM, PA designed the study; YL, MM, HG, MK, ARM, AM performed the analysis; YL, MM, HG, MK, ARM, WM, PA wrote the manuscript.

## 8 Acknowledgments

The authors would like to thank Clark Gao, Lori Jean Van Orden, Martine Meyer, and Simone Krupka for their help and discussions in shaping this work.

