## Supplementary material for "Highly Predictive Transdiagnostic Features Shared across Schizophrenia, Bipolar Disorder, and ADHD Identified Using a Machine Learning Based Approach"

**Supplementary Figures**

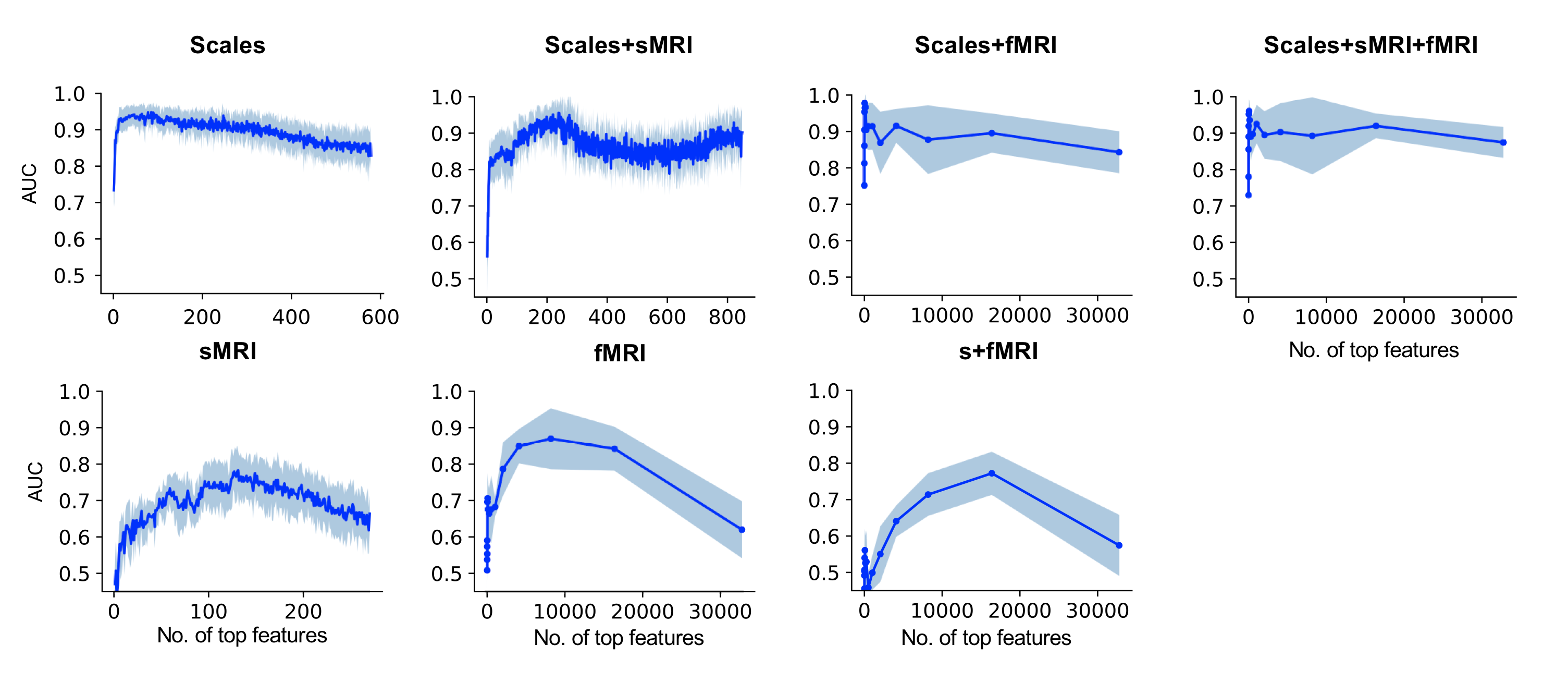

**Supplementary Figure 1.** Results of the feature importance-guided sequential model selection for the HC vs. All Patients transdiagnostic model. Performances of the truncated models (AUC) are plotted as a function of the number of top features used as input. The blue line represents the mean AUC across 10 iterations of the sequential model selection procedure and the shaded region represents the mean ± 1 standard deviation.

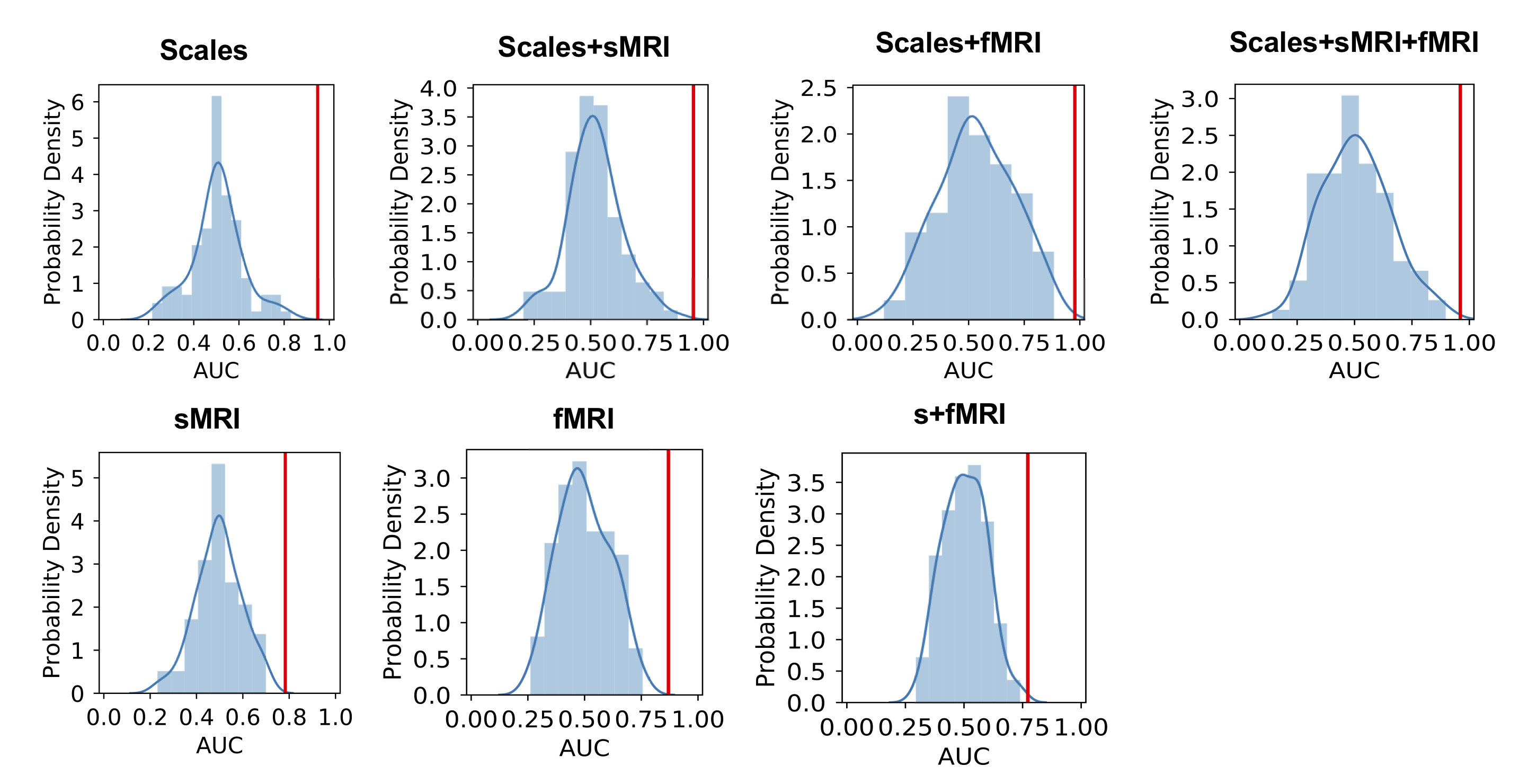

**Supplementary Figure 2.** Permutation test results on performances of the best truncated transdiagnostic models. The histogram represents the empirical null distribution constructed by evaluating model performances on 100 iterations of label shuffled data. The red vertical line marks the actual AUC obtained from the original data. All transdiagnostic models performed significantly higher than chance level (AUC=0.5; p < 0.01, FDR corrected).

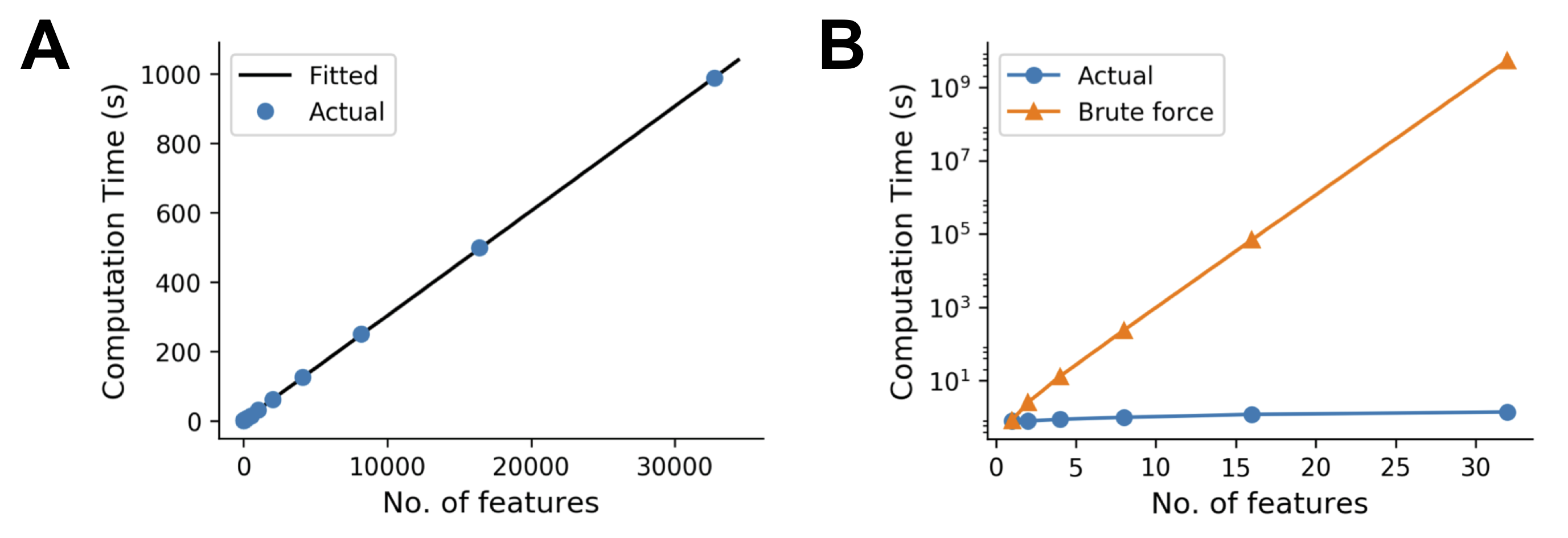

**Supplementary Figure 3.** Time complexity of the importance-guided sequential model selection procedure. A) The computation time (median across 3 implementations) grew linearly as the number of features increases in our importance-guided forward model selection procedure. The blue dots represent the actual measured data points, whereas the black line represents the fitted regression line (slope: 0.03; intercept: 0.80). B) Significantly reduced computation time was achieved via our sequential model selection procedure (blue) compared to the estimated time complexity of a brute force feature selection procedure where all combinations of features are evaluated (orange).

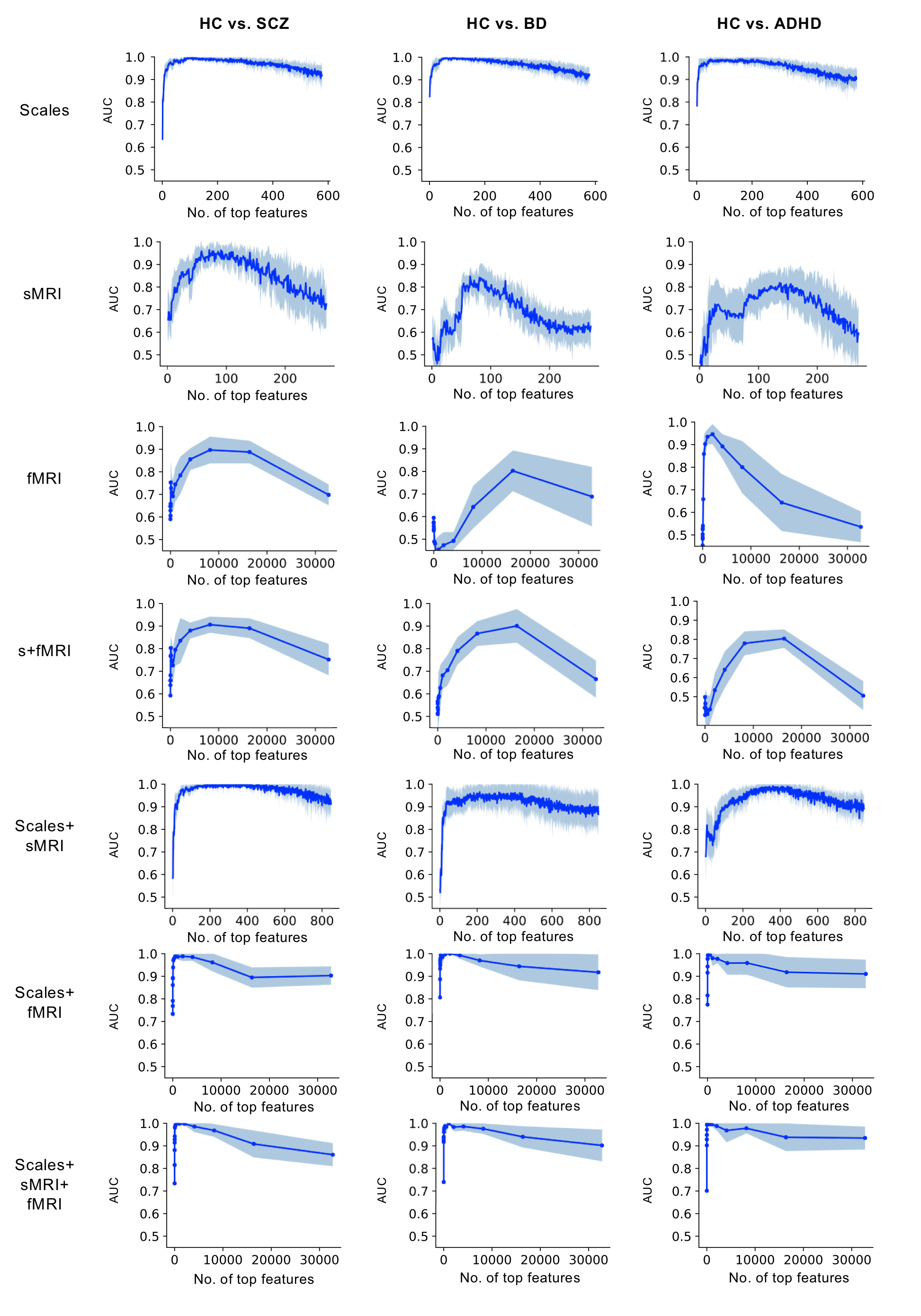

**Supplementary Figure 4.** Results of the feature importance-guided sequential model selection for the 3 HC vs. a single patient category models. The AUCs from the truncated models are plotted as a function of the number of top features used as inputs.

**Supplementary Tables**

**Supplementary Table 1. Scores from the 13 Self-Reported Instruments**

| **Sum/Subscores** | **HC** | | **SCZ** | | **BD** | | **ADHD** | |
| --- | --- | --- | --- | --- | --- | --- | --- | --- |
|  | **Mean** | **SD** | **Mean** | **SD** | **Mean** | **SD** | **Mean** | **SD** |
| Chapman Social Anhedonia Total | 9.68 | 6.67 | 14.94 | 6.03 | 16.35 | 7.29 | 13.47 | 8.22 |
| Chapman Physical Anhedonia Total | 11.55 | 6.46 | 16.72 | 6.96 | 15.90 | 8.72 | 12.98 | 7.49 |
| Chapman Hypomanic Personality Total | 16.64 | 7.62 | 19.30 | 8.78 | 25.18 | 11.32 | 25.09 | 7.27 |
| Chapman Perceptual Aberration Total | 2.03 | 2.41 | 9.12 | 8.02 | 5.14 | 4.51 | 4.33 | 4.39 |
| Hopkins Symptom Checklist |  |  |  |  |  |  |  |  |
| Global Severity Score | 0.32 | 0.24 | 0.84 | 0.52 | 0.87 | 0.55 | 0.68 | 0.39 |
| Anxiety Sub-score | 0.20 | 0.27 | 0.84 | 0.72 | 0.74 | 0.62 | 0.47 | 0.45 |
| Depression Sub-score | 0.37 | 0.35 | 0.83 | 0.63 | 0.96 | 0.63 | 0.69 | 0.50 |
| Interpersonal Sensitivity Sub-score | 0.39 | 0.35 | 0.96 | 0.68 | 1.09 | 0.72 | 0.76 | 0.51 |
| Obsessive Compulsive Sub-score | 0.49 | 0.42 | 1.06 | 0.64 | 1.16 | 0.72 | 1.24 | 0.70 |
| Somatization Sub-score | 0.21 | 0.23 | 0.66 | 0.53 | 0.64 | 0.60 | 0.41 | 0.34 |
| Temperament and Character Inventory |  |  |  |  |  |  |  |  |
| Harm Avoidance Sub-score | 11.11 | 6.20 | 16.06 | 7.48 | 17.76 | 9.02 | 12.98 | 6.80 |
| Novelty Seeking Sub-score | 19.36 | 6.07 | 18.50 | 4.94 | 24.45 | 8.02 | 25.33 | 4.88 |
| Persistence Sub-score | 24.55 | 6.89 | 21.46 | 6.70 | 19.00 | 9.47 | 21.51 | 7.74 |
| Reward Dependence Sub-score | 16.08 | 4.07 | 14.36 | 3.91 | 13.63 | 5.09 | 15.02 | 4.65 |
| Adult ADHD Self-Report Scale Total | 8.67 | 2.90 | 9.86 | 4.52 | 13.20 | 4.89 | 15.49 | 3.91 |
| Barratt Impulsiveness Scale |  |  |  |  |  |  |  |  |
| Attentional Sub-score | 14.38 | 3.45 | 17.28 | 4.66 | 19.65 | 5.10 | 21.67 | 4.03 |
| Motor Sub-score | 21.66 | 3.84 | 22.96 | 4.79 | 25.53 | 5.52 | 26.60 | 4.17 |
| Non-planning Sub-score | 22.88 | 4.40 | 26.26 | 5.28 | 29.31 | 5.58 | 28.53 | 4.45 |
| Dickman Impulsivity Scale |  |  |  |  |  |  |  |  |
| Functional Impulsivity Total | 6.59 | 2.61 | 5.40 | 2.65 | 5.88 | 3.25 | 6.84 | 2.86 |
| Dysfunctional Impulsivity Total | 1.95 | 2.37 | 4.18 | 3.32 | 6.35 | 4.43 | 4.98 | 3.15 |
| Multidimensional Personality Q Total | 18.12 | 4.91 | 15.94 | 4.85 | 10.18 | 7.37 | 10.95 | 5.06 |
| Eysenck’s Impulsivity Inventory |  |  |  |  |  |  |  |  |
| Impulsiveness Sub-score | 6.21 | 2.82 | 9.14 | 3.43 | 9.82 | 4.29 | 9.47 | 3.43 |
| Venturesomeness Sub-score | 8.78 | 2.48 | 8.02 | 2.84 | 8.00 | 2.53 | 9.44 | 1.82 |
| Empathy Sub-score | 10.53 | 3.09 | 10.84 | 3.36 | 11.18 | 3.40 | 11.51 | 2.86 |
| Scale for Bipolar II Risk |  |  |  |  |  |  |  |  |
| Mood Liability Sub-score | 2.08 | 1.68 | 3.96 | 2.66 | 5.67 | 2.80 | 3.98 | 2.44 |
| Daydreaming Sub-score | 2.92 | 1.76 | 3.02 | 1.98 | 3.82 | 1.73 | 3.92 | 1.38 |
| Energy-Activity Sub-score | 3.27 | 2.10 | 3.96 | 2.07 | 4.16 | 2.54 | 3.86 | 2.12 |
| Social Anxiety Sub-score | 2.90 | 1.76 | 3.60 | 1.71 | 3.73 | 1.85 | 3.09 | 1.78 |
| Summary Score | 11.17 | 4.23 | 14.54 | 5.72 | 17.39 | 6.45 | 14.84 | 4.01 |
| Golden and Meehl’s 7 MMPI Items | 2.40 | 1.24 | 3.4 | 1.81 | 4.04 | 1.59 | 3.09 | 1.36 |

**Supplementary Table 2. Effect Sizes between HC and Patients based on sum scores/subscores**

| Sum/Subscores | Cohen's d |
| --- | --- |
| Hopkins Global Severity | 1.21 |
| Barratt Impulsiveness Attentional | 1.17 |
| Hopkins Obsessive Compulsive | 1.14 |
| Bipolar II Mood | 1.07 |
| Barratt Impulsiveness Nonplanning | 1.05 |
| Hopkins Interpersonal Sensitivity | 1.03 |
| Dickman Dysfunctional Total | 1.01 |
| Hopkins Anxiety | 0.99 |
| Multidimensional Personality Control | 0.99 |
| Eysenck Impulsiveness Score | 0.98 |
| Adult ADHD Total | 0.98 |
| Hopkins Depression | 0.92 |
| Hopkins Somatization | 0.91 |
| Bipolar II Sumscore | 0.88 |
| Chapman Perceptual Aberration Total | 0.88 |
| Golden and Meehl's 7 MMPI Items Total | 0.77 |
| Chapman Social Anhedonia Total | 0.76 |
| Chapman Hypomanic Personality Total | 0.74 |
| Barratt Impulsiveness Motor | 0.73 |
| TCI Harm Avoidance | 0.64 |
| TCI Persistence | 0.52 |
| Chapman Physical Anhedonia Total | 0.52 |
| TCI Novelty Seeking | 0.50 |
| TCI Reward Dependence | 0.41 |
| Bipollar II Daydreaming | 0.37 |
| Bipollar II Energy | 0.34 |
| Bipolar II Anxiety | 0.33 |
| Dickman Functional Total | 0.21 |
| Eysenck Empathy Score | 0.20 |
| Eysenck Venturesomeness Score | 0.13 |
| Sum of Top 85 Transdiagnostic Items | 2.85 |

| **Supplementary Table 3. Performances of the best truncated models for HC vs. each patient category*** | | | | | | |  |
| --- | --- | --- | --- | --- | --- | --- | --- |
| ***HC vs. SCZ:*** |  |  |  |  |  |  |  |
|  | Scales | sMRI | fMRI | s+fMRI | Scales+sMRI | Scales+fMRI | Scales+s+fMRI |
| AUC | 0.997(0.007) | 0.964(0.038) | 0.897(0.060) | 0.906(0.036) | 0.999(0.003) | 0.989(0.021) | 1.000(0.001) |
| Accuracy | 0.969(0.023) | 0.885(0.030) | 0.868(0.044) | 0.865(0.038) | 0.877(0.038) | 0.958(0.048) | 0.903(0.080) |
| Sensitivity | 0.930(0.078) | 0.517(0.117) | 0.870(0.119) | 0.910(0.164) | 0.467(0.163) | 0.920(0.125) | 0.700(0.249) |
| Specificity | 0.985(0.019) | 0.995(0.015) | 0.867(0.079) | 0.843(0.083) | 1.000(0.000) | 0.976(0.044) | 1.000(0.000) |
| No. of features | 112 | 71 | 8192 | 8192 | 333 | 2048 | 512 |
| ***HC vs. BD:*** |  |  |  |  |  |  |  |
|  | Scales | sMRI | fMRI | s+fMRI | Scales+sMRI | Scales+fMRI | Scales+s+fMRI |
| AUC | 0.998(0.003) | 0.847(0.038) | 0.803(0.090) | 0.901(0.074) | 0.966(0.042) | 1.000(0.000) | 0.999(0.002) |
| Accuracy | 0.978(0.027) | 0.810(0.032) | 0.769(0.095) | 0.872(0.093) | 0.883(0.064) | 0.979(0.032) | 0.966(0.022) |
| Sensitivity | 0.930(0.090) | 0.567(0.116) | 0.762(0.142) | 0.838(0.210) | 0.667(0.243) | 0.925(0.115) | 0.912(0.080) |
| Specificity | 0.996(0.012) | 0.920(0.046) | 0.771(0.144) | 0.886(0.125) | 0.980(0.033) | 1.000(0.000) | 0.986(0.030) |
| No. of features | 64 | 65 | 16384 | 16384 | 213 | 2048 | 1024 |
| ***HC vs. ADHD:*** |  |  |  |  |  |  |  |
|  | Scales | sMRI | fMRI | s+fMRI | Scales+sMRI | Scales+fMRI | Scales+s+fMRI |
| AUC | 0.991(0.011) | 0.819(0.079) | 0.946(0.045) | 0.804(0.049) | 0.991(0.009) | 0.996(0.009) | 0.999(0.002) |
| Accuracy | 0.940(0.027) | 0.752(0.041) | 0.890(0.069) | 0.769(0.062) | 0.822(0.036) | 0.876(0.058) | 0.910(0.047) |
| Sensitivity | 0.844(0.102) | 0.171(0.057) | 0.725(0.289) | 0.875(0.125) | 0.314(0.140) | 0.550(0.211) | 0.675(0.170) |
| Specificity | 0.973(0.046) | 0.955(0.052) | 0.952(0.074) | 0.729(0.102) | 1.000(0.000) | 1.000(0.000) | 1.000(0.000) |
| No. of features | 165 | 136 | 2048 | 16384 | 360 | 128 | 128 |

* The mean performance measures across 10 implementations are reported here with the standard deviation shown in parentheses

| **Supplementary Table 4. Grouping of the top 20 items from the HC vs. Patients transdiagnostic classifier** | |
| --- | --- |
| *Mood dysregulation:* | |
| BipolarII 1 | Mood changes without knowing why |
| Chaphypo 21 (-) | Moods do not seem to fluctuate more than most people |
| *Psychomotor agitation:* | |
| Chaphypo 8 | So restless; impossible to sit still |
| *Inattention:* |  |
| Hopkins 55 | Trouble concentrating |
| ASRS | Trouble wrapping up final details of a project |
| TCI 35t | Difficult to keep the same interests because attention shifts |
| *Social Anhedonia:* | |
| Chapsoc 8 (-) | Have more fun doing things with other people |
| Chapsoc 40 | Prefer company of pets to people |
| BipolarII 26 (-) | Never get all that I need from people |
| *Apathy:* |  |
| TCI 72p (-) | I love to excel at everything I do |
| Dickman 5 (-) | Have many hobbies |
| TCI 228p (-) | Work extra hard to correct mistakes |
| *Neuroticism:* |  |
| Eysenck 18 | Friends upset affect you? |
| Hopkins36 | Feel others do not understand you or are unsympathetic |
| *Impulsivity:* |  |
| Eysenck 31 | Need a lot of self-control to keep out of trouble |
| TCI 61t (-) | Think about things for a long time before decision |
| TCI 13t | Do things based on how I feel at the moment |
| *Extraversion:* |  |
| Chaphypo 34 | So many fields I could succeed in; a shame to have to pick |
| TCI 68t (-) | Keep problems to oneself |
| *Religion:* |  |
| Dickman 27 | Religion is very important |
| (-) denotes a negative model weight; the corresponding item is reversed during grouping | |

| **Supplementary Table 5. Grouping of the top 20 items from the HC vs. SCZ classifier** | |
| --- | --- |
| *Mood dysregulation:* | |
| BipolarII 1 | Mood changes without knowing why |
| *Psychomotor agitation:* | |
| Chaphypo 8 | So restless; impossible to sit still |
| *Perceptual aberration:* | |
| Chapper 11 | Body is abnormal |
| Chapper 35 | Heightened awareness of sights and sounds; cannot shut them out |
| *Apathy:* |  |
| BipolarII 19 (-) | Like to indulge in a reverie |
| Dickman 29 (-) | Have more curiosity than most people |
| TCI 189p | Like to go slow in starting work, even easy to do |
| Chaphypo 26 (-) | Would make a good actor; can play many roles convincingly |
| *Social anhedonia:* | |
| Chapsoc 23 (-) | Expect me to spend more time talking with people |
| TCI 240t (-) | Talk about people behind their backs |
| BipolarII 26 (-) | Never get all that I need from people |
| Chaphypo 13 (-) | People come to me for a clever idea |
| Eysenck 53 (-) | Can imagine what it is like to be lonely |
| *Physical Anhedonia:* | |
| Chapphy 29 (-) | Smell of fresh bread make me hungry |
| *Neuroticism:* |  |
| BipolarII 22 (-) | Inclined to think about myself |
| TCI 110t (-) | Tell a funny story or to play a joke on someone |
| *Distress:* |  |
| Golden 1 | I have not lived the right kind of life |
| Hopkins 22 | Feeling of being trapped or caught |
| Hopkins 33 | Feeling fearful |
| TCI 129t | Feel tense and worried in unfamiliar situations |
| (-) denotes a negative model weight; the corresponding item is reversed during grouping | |

| **Supplementary Table 6. Grouping of the top 20 items from the HC vs. BD classifier** | |
| --- | --- |
| *Mood dysregulation:* | |
| Chaphypo 21 (-) | Moods do not seem to fluctuate more than most people |
| BipolarII 1 | Mood changes without knowing why |
| Chaphypo 17 (-) | I am usually in an average sort of mood |
| *Social anhedonia:* |  |
| BipolarII 26 (-) | Never get all that I need from people |
| Chapsoc 36 (-) | Would much rather be with others than be alone |
| Chapsoc 40 | Prefer accompany of pets to people |
| *Physical anhedonia:* | |
| Chapphy 13 | Sound of rustling leaves never pleased me |
| Chapphy 2 (-) | Tried to eat slowly when eating favorite food |
| *Impulsivity:* |  |
| Eysenck 31 | Need a lot of self-control to keep out of trouble |
| Eysenck 44 (-) | Consider all pros and cons before making a decision |
| TCI 34t (-) | Like to be very organized and set up rules |
| *Increased energy:* |  |
| BipolarII 17 | Experienced periods that sleep isn't necessary for days |
| *Apathy:* |  |
| Chaphypo 19 (-) | Have a wide range of interests |
| Dickman 5 (-) | Have many hobbies |
| BipolarII 15 (-) | Quick in actions |
| *Religion:* |  |
| Dickman 27 | Religion is very important in my life |
| *Inattention:* |  |
| ASRS | Trouble wrapping up final details of a project |
| *Extraversion:* |  |
| Chaphypo 1 (-) | Consider oneself to be an average kind of person |
| *Distress:* |  |
| TCI 129t | Feel tense and worried in unfamiliar situations |
| BipolarII 5 | Future looks very dark |
| (-) denotes a negative model weight; the corresponding item is reversed during grouping | |

| **Supplementary Table 7. Grouping of the top 20 items from the HC vs. ADHD classifier** | |
| --- | --- |
| *Mood dysregulation:* | |
| BipolarII 1 | Mood often changes without knowing why |
| Chaphypo 17 (-) | I am usually in an average sort of mood |
| *Agitation:* |  |
| Chaphypo 8 | So restless; impossible to sit still |
| Barratt 28 | Restless at lectures or talks |
| Chaphypo 32 | Considered to be a "hyper" person |
| *Inattention:* |  |
| Hopkins 55 | Trouble concentrating |
| Barratt 9 (-) | Concentrate easily |
| Barratt 26 | Have outside thoughts when thinking |
| *Extraversion:* |  |
| Chaphypo 26 | Would make a good actor; can play many roles convincingly |
| TCI 204t (-) | Not very good at talking my way out of trouble |
| TCI 52t (-) | Much better as a listener than as a talker |
| *Physical anhedonia:* | |
| Chapphy 7 (-) | Taste of food has always been important |
| *Social anhedonia:* |  |
| Chapsoc 8 (-) | Have more fun doing things with other people |
| Chapsoc 17 | Prefer activities that do not involve other people |
| Chapsoc 24 (-) | Feel pleased and gratified learn more about emotional life of friends |
| *Indecision:* |  |
| Dickman 30 (-) | Like sports/games that have to choose moves very quickly |
| *Impulsivity:* |  |
| TCI 61t (-) | Think about things for a long time before decision |
| Dickman 28 | Get into trouble because I don't think before act |
| MPQ 12 | Stop one thing before completing it and start another |
| TCI 82t (-) | Think about all the facts before making a decision |
| (-) denotes a negative model weight; the corresponding item is reversed during grouping | |
